# *In silico* target discovery for aromatic amino acid production in *Escherichia coli*

**DOI:** 10.1101/2024.11.25.625200

**Authors:** André Fonseca, Leslie Avendãno, Sofia Ferreira, Isabel Rocha

## Abstract

Aromatic amino acids (AAA) and derived compounds are rising on global market demand, which leads to the need of better production means. These compounds are usually produced by means of microbial cell factories. However, bacterial metabolism is complex and highly regulated, which complicates the discovery of new higher yield strains, optimized for the production of these compounds. Dynamic modelling has the capability to solve this problem by predicting metabolic behavior and helping to find targets for strain design. In this paper, a new *Escherichia coli* dynamic model is presented. Evolutionary algorithm processes were used in order to find targets for metabolic engineering strategies for AAA increased production. The solutions obtained are compatible with strategies found in literature, while at the same time presenting new possible targets, not yet reported in literature.

## 1. Introduction

Amino acid global demand has been steadily increasing in the last decades, reaching a market size valued at 21 billion dollars (USD) in 2019, which is estimated to keep increasing in the next few years^1^. Aromatic amino acids (AAA), in particular, are of great interest due to also being the gateway to other high-added value aromatic compounds. AAA and their derived compounds have various applications, such as food and feed additives, pharmaceuticals, cosmetics, dietary supplements and others^2–6^. Currently, AAA production is mainly obtained by microbial processes and there are industrial strains used solely for the production of each individual AAA: phenylalanine (L-Phe), tryptophan (L-Trp) and tyrosine (L-Tyr)^6–9^. However, the constant market growth for AAA makes current production insufficient to keep up with ever increasing demands, implying a need for better solutions, such as higher yield strains. One of the biggest hurdles for higher efficiency strains lies in the high complexity of microbial metabolism, both due to its size and its complex regulation^10–12^. Mathematical models can help to better understand these intricacies by providing predictions of metabolic behavior for specific phenotypes and environmental conditions^13^. There are two main approaches for mathematical modeling of biological systems: constraint based modeling and dynamic modeling. These approaches have different assumptions, with the first assuming that metabolites are in a pseudo steady state, with constant concentration; the second approach assumes a less rigid position, acknowledging the changes in metabolite concentration that naturally occur over time^14,15^.

Metabolic processes are dependent on several factors such as pH, temperature, metabolite concentration, etc. These factors are naturally fluctuating, meaning that the temporal behavior of metabolic processes such as enzymatic reactions are not static. As such, dynamic (or kinetic models as they are also called) have the potential to accurately predict biological systems, by taking these factors into account^14,16,17^. In these kinds of models, mathematical expressions are used to represent the rates at which reactions operate over time. These expressions are joined into a set of ordinary differential equations (ODEs), which represents the entire modeled system. The rates composing the ODEs can be mechanistic in nature, usually obtained from *in vitro* studies, but they can also be stochastic, obtained by fitting experimental data into approximate mathematical expressions, if there is a lack of kinetic information^13,18^. Due to their level of detail, dynamic models can predict and quantify reaction fluxes, metabolites, and even inhibitory regulation, making them great tools for strain target discovery^19^.

In this paper, we intend to demonstrate a new *Escherichia coli* dynamic model, and its capability to accurately simulate both wild type and mutant strains. We also intend to use this model to predict targets for metabolic engineering, with the objective of creating strains capable of overproducing each of the AAA: phenylalanine, tyrosine, and tryptophan.

## 2. Results and discussion

### 2.1. Model description

*Escherichia coli* has one of the best characterized metabolic networks in all of biology, due to its industrial and scientific importance, being regarded as the prime microbial model^10,20^. *E. coli*’s central carbon metabolism (CCM), the main hub for cell biosynthesis and catabolism, has been extensively studied, with several dynamic models being made to simulate and represent its activity^12,16,21–23^. However, there is still missing information regarding *E. coli*’s metabolism, particularly enzyme kinetic information. This severely hampers the construction of dynamic models, as they rely on enzyme kinetics for building the ODEs that represent metabolic processes. Lack of knowledge is one of the greatest limitations to the creation of bigger, more accurate dynamic models^16^. In fact, up to this point there were no reported dynamic models that represented an extended version of the CCM, including the shikimate pathway and the three AAA production pathways. Therefore, instead of “reinventing the wheel” many reactions and pathways, including enzyme kinetics, that were used to construct the model presented in this paper were either derived or directly taken from existing dynamic models in the literature, as described in the Results section, with the exception of the shikimate and AAA biosynthesis pathways. Being constructed from scratch, and with them being a major part of this work, great care was taken to ensure that kinetics and end product feedback inhibition were properly represented in these pathways.

In *E. coli*, the first reaction of the shikimate pathway (DDPA) is catalyzed by three isozymes, each one feedback inhibited by one of the AAA: phenylalanine, tryptophan, or tyrosine. These isozymes are responsible for the conversion of erythrose-4-phosphate (E4P) and phosphoenolpyruvate (PEP) into 2-dehydro-3-deoxy-D-arabino-heptonate 7-phosphate (2dda7p). The remaining six reactions of the shikimate pathway aren’t regulated by feedback inhibition, although some product inhibition can be found. The last reaction of this pathway produces chorismate, which is a common precursor for all three AAA biosynthetic pathways. In tryptophan’s case, two L-Trp feedback inhibited reactions (ANS and ANPRT) catalyze chorismate into N-(5-Phospho-D-ribosyl)anthranilate, which will be converted into tryptophan after four more reactions, none of which are feedback inhibited by tryptophan. In phenylalanine’s case, a L-Phe feedback inhibited bifunctional enzyme is responsible for the two-step conversion of chorismate into prephenate and then into phenylpyruvate. This metabolite is then converted to phenylalanine by a transaminase reaction. Tyrosine has a similar pathway to phenylalanine, with a L-Tyr feedback inhibited bifunctional enzyme responsible for the two-step conversion of chorismate into prephenate and then into 3-(4-hydroxyphenyl)pyruvate. The metabolite is then converted to tyrosine by a transaminase reaction. The enzyme kinetics used for the construction of the shikimate pathway and AAA production can be seen in Table 1. A schematic of the entire dynamic model, including the AAA regulatory processes, can be seen in Figure 1.

**Table 1.**
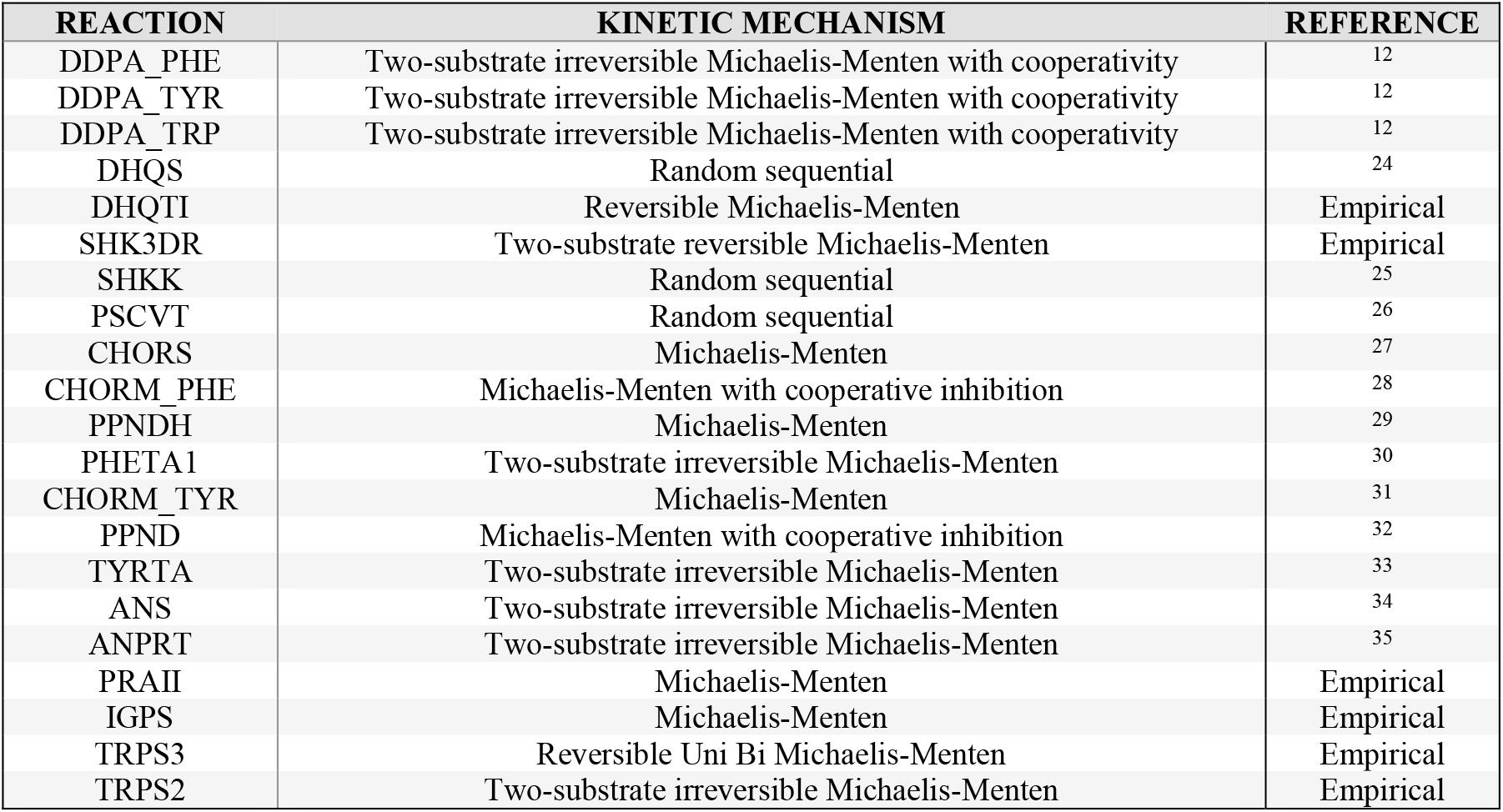
Kinetic mechanism of the reactions from the shikimate pathway and the AAA pathways. Unlike reactions from other pathways in the model, these were based on kinetic data found on the literature, instead of existing kinetic models, with the exception of the DDPA isozymes (DDPA_PHE, DDPA_TYR and DDPA_TRP) which were a mix of both. The kinetic mechanisms of some reactions are empirical due to insufficient kinetic data.

**Figure 1.**
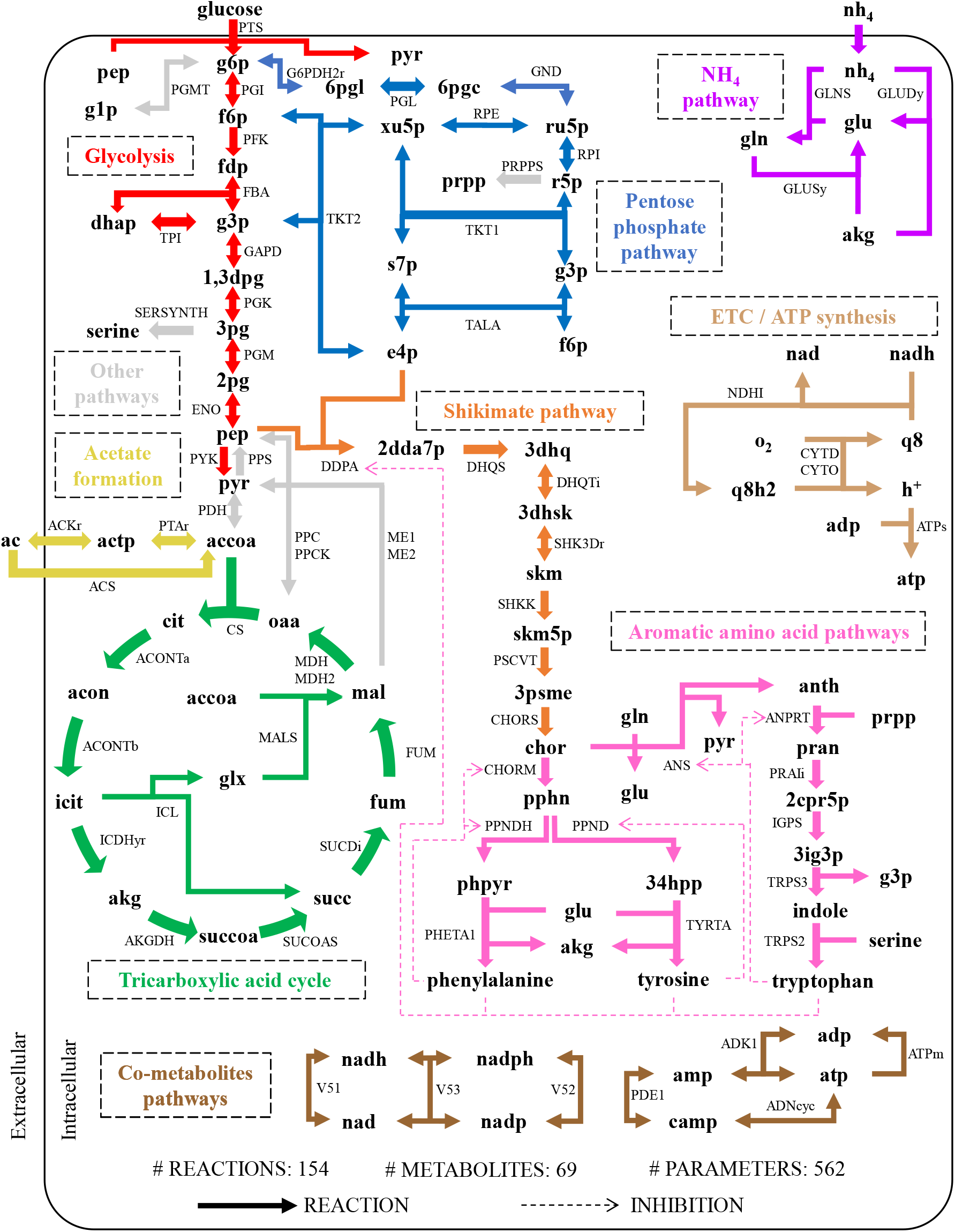
Scheme showing the structure of the dynamic model. Arrows correspond to reactions. Dashed arrows correspond to inhibition. There are more inhibitory processes in the model, but only the AAA ones are shown for an easier to read scheme. Each pathway is represented by a distinct color: red – glycolysis, blue – pentose phosphate pathway, green -tricarboxylic acid cycle, yellow – acetate formation, purple – ammonia intake, orange – shikimate pathway, pink – AAA formation pathways, light brown – electron transport chain and ATP synthesis, dark brown – co-metabolite maintenance and pathway reactions, and grey – other pathways.

### 2.2. Simulating WT and mutant strains

Simulations were performed to test the dynamic model against experimentally obtained data, found in literature. For experimental data we used the steady state fluxes obtained by Ishii et al. (2007)^36^, in which *E. coli* was grown in glucose limited chemostat cultures with glucose concentration in the feed set to 4 g/L. Also, experimental data shows steady state fluxes of WT cultures grown with different dilution rates: 0.1 h^-1^, 0.4 h^-1^, 0.5 h^-1^ and 0.7 h^-1^, as well as fluxes for several single gene knockout (KO) mutant strains, all grown with a dilution rate of 0.2 h^-1^. These conditions were considered when simulating *in silico*. The sets of fluxes, both simulated and experimental, were thus normalized to glucose intake by the phosphotransferase reaction and compared. Similarly, simulations with a stoichiometric model from the literature (iJO1366^37^) were also used, with its fluxes normalized and compared to experimental data as well. Since there are no other dynamic models, to our knowledge, that can simulate the shikimate and AAA pathways a stoichiometric one was chosen for comparison. We intended to analyze the differences between both types of models in regard to the simulation of mutant strains and growth parameters. The iJO1366 model was chosen as it is one of the most complete and accurate metabolic reconstructions of *E. coli*, being one of the main references for phenotypic predictions of this organism. To represent the fit between experimental and model data we used a linear regression between experimental and simulated data and calculated the square of the regression coefficient (R^2^).

The dynamic model appears to be better at predicting data from higher dilution rates. In fact, the fit for the lowest experimental rate of 0.1 h^-1^ is the worst R^2^ value achieved by this model (0.77). The other dilution rates simulations present much higher prediction power, with the 0.5 h^-1^ rate having a R^2^ value of 0.951, one of the highest achieved by this model and highest among the WT simulations. In contrast, the stoichiometric model shows similar levels of prediction across all dilution rates, even though they aren’t as high as the dynamic model simulations. Similar to the dynamic model, iJO1366 also seems to better predict data from the dilution rate of 0.5 h^-1^, having an R^2^ value of 0.908, the highest among the WT simulations. The regressions for the different dilution rates simulations are shown in Figure 2.

**Figure 2.**
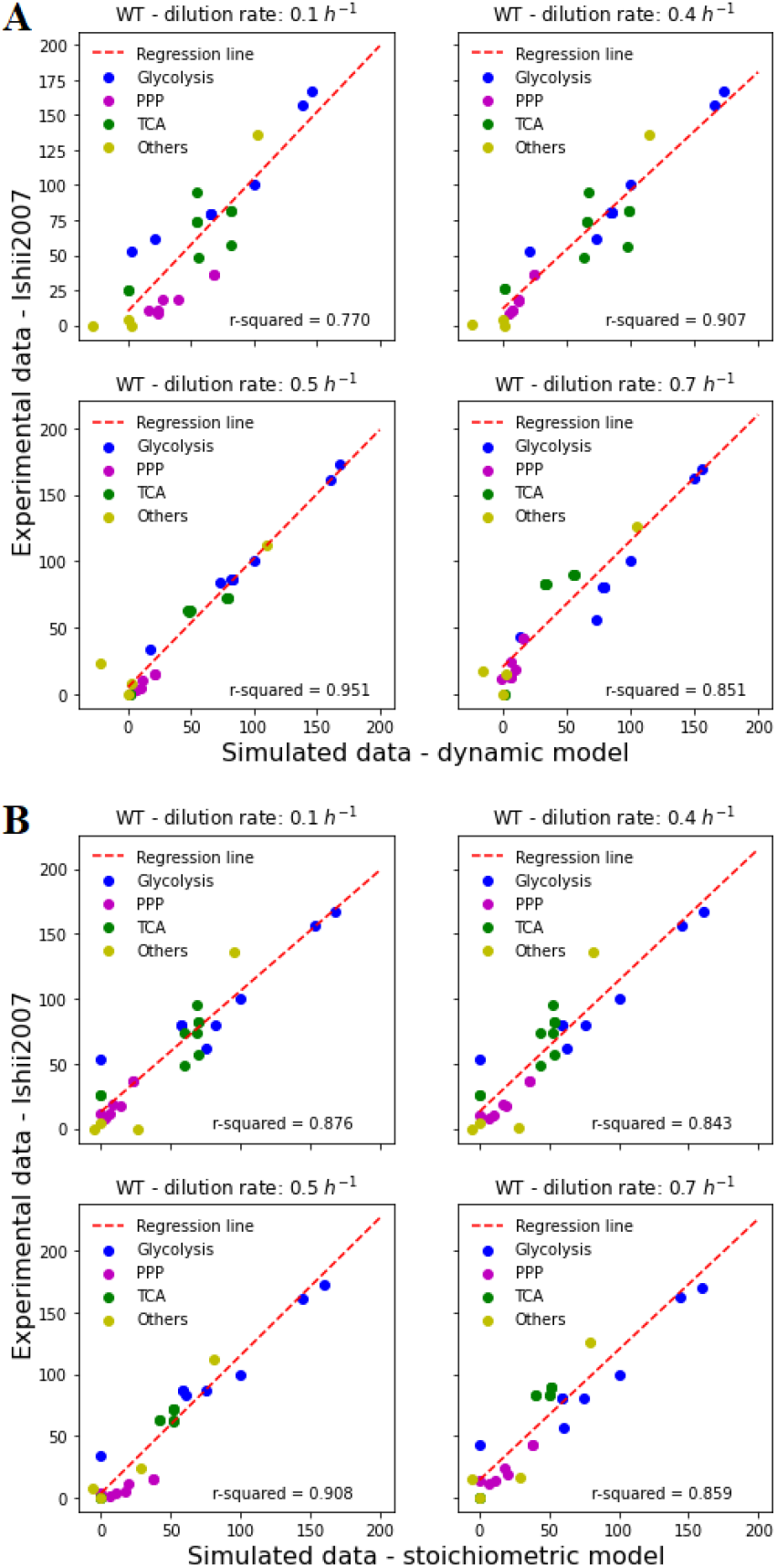
Linear regressions of dynamic (A) and stoichiometric (B) fluxes with experimental data (Ishii et al. 2007) for different dilution rates. Reaction fluxes are divided by color according to the pathway attributed to them: blue – glycolysis; purple – pentose phosphate pathway (PPP); green – tricarboxylic acid cycle (TCA); yellow – other pathways. Values of squared regression coefficient (r-squared) are written below each image.

The single gene KO strain simulations by the dynamic model show great fit between simulated and experimental data, with seven out of ten KO simulations having an R^2^ value above 0.9, with the highest being the rpiB gene KO with 0.966 and the lowest being the pfkA gene KO with 0.807. In contrast, the stoichiometric model presents a worse fit to experimental data, with only three out of 10 KO simulations having an R^2^ value above 0.9. For this model, the highest R^2^ value was 0.942 with the ppsA KO simulation and the lowest was 0.721 with the tktA KO simulation. These values seem to indicate the dynamic model as a valid predictor of WT and mutant strain phenotype. The regressions for the different single gene KO simulations are shown in Figure 3.

**Figure 3.**
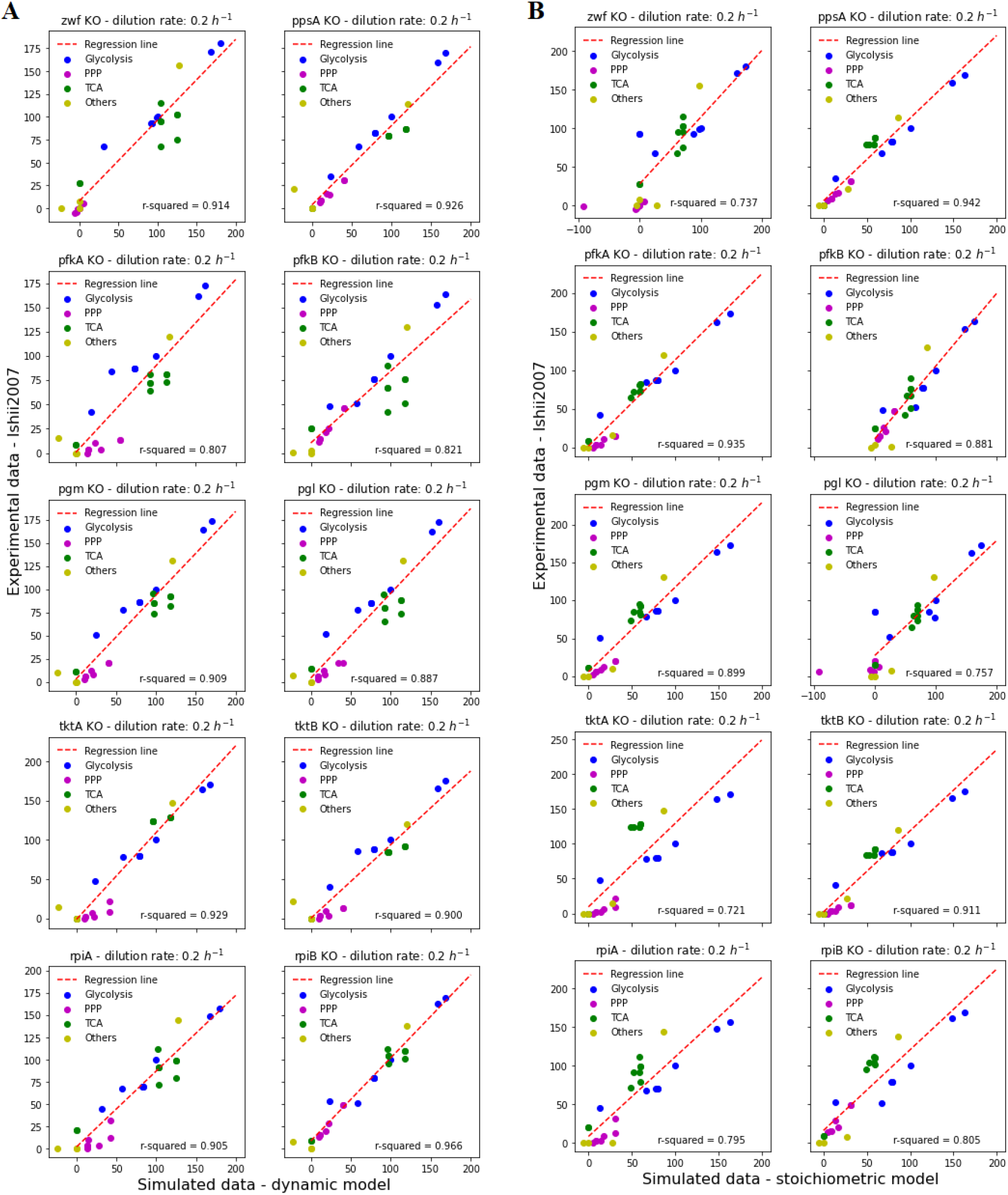
Linear regressions of dynamic (A) and stoichiometric (B) fluxes with experimental data (Ishii et al. 2007) for different single gene knockout strains. Reaction fluxes are divided by color according to the pathway attributed to them: blue – glycolysis; purple – pentose phosphate pathway (PPP); green – tricarboxylic acid cycle (TCA); yellow – other pathways. Values of squared regression coefficient (r-squared) are written below each image.

**Figure 4.**
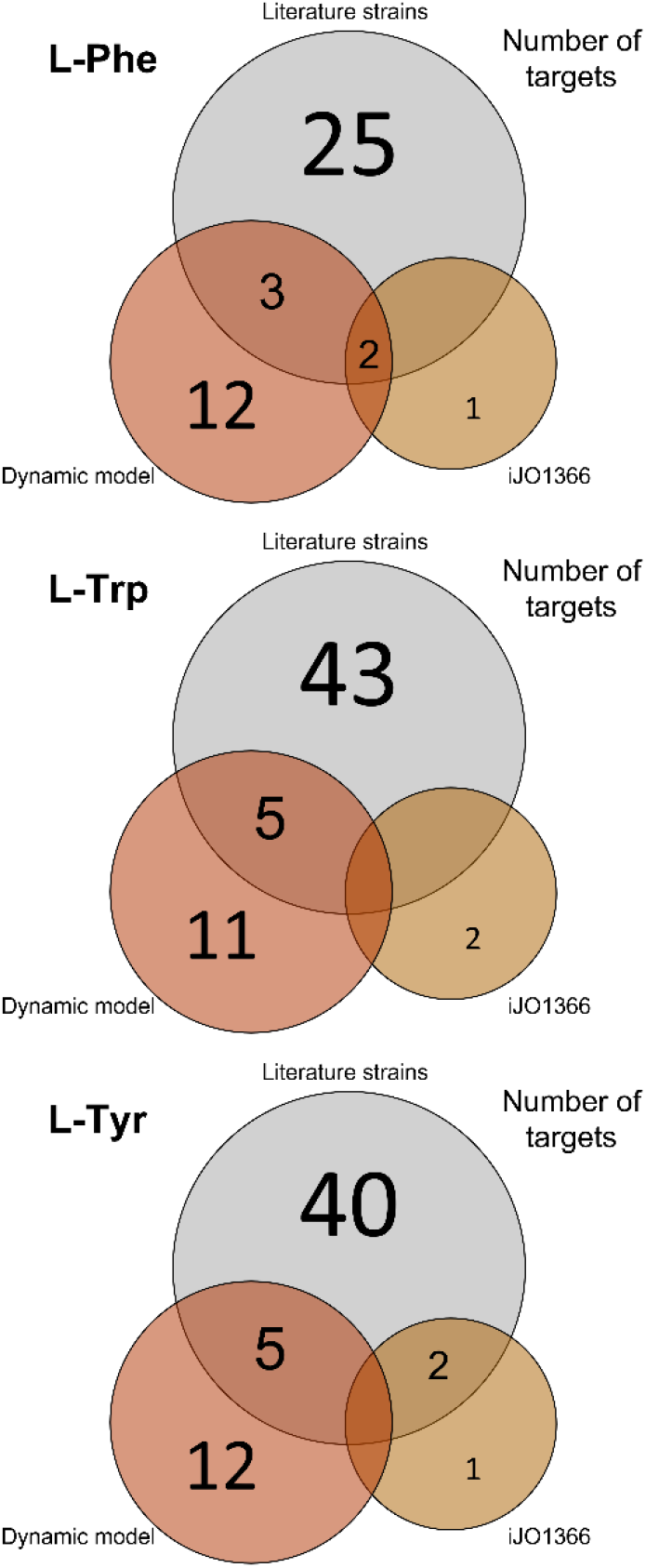
Common targets found in aromatic amino acids overproducing strains from three different sources presented as Venn diagrams: literature, strain target optimization of the dynamic model presented in this work and also of the stoichiometric model iJO1366. Top left is for phenylalanine (L-Phe), top right for tyrosine (L-Tyr) and bottom left for tryptophan (L-Trp).

**Figure 5.**
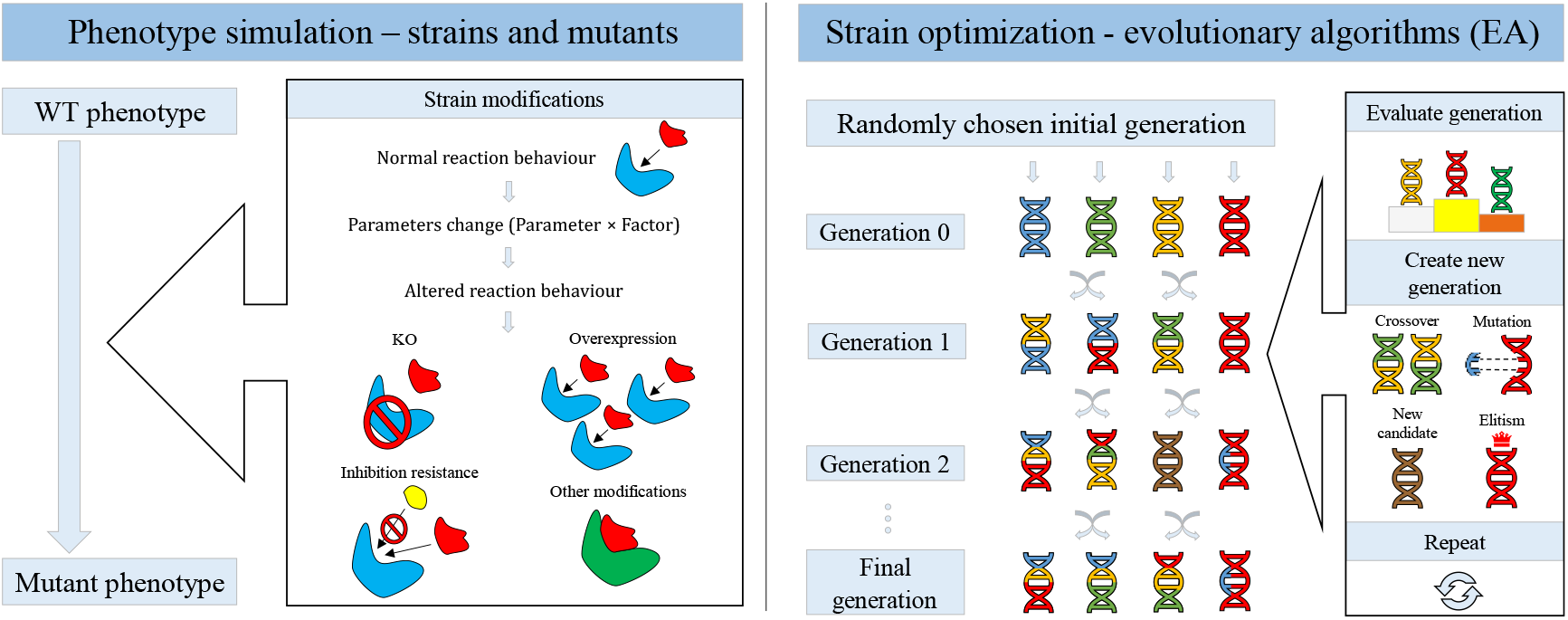
Schematic representation of phenotype simulation (left) and strain optimization (right) with optimModels. The mutant phenotype is dependent on the parameter(s) and factor(s) chosen. The objective function used to determine candidates’ fitness and evaluate each generation was the steady state concentration of either Phenylalanine, Tryptophan or Tyrosine. Two hundred generations was the maximum limit for each optimization.

Overall, the dynamic model simulations present a higher fit to experimental values, represented by a higher average R^2^ value of 0.889 opposed to the stoichiometric model’s R^2^ value of 0.848. This implies a tendency for better prediction ability by the dynamic model. Unfortunately, there is a lack of experimental data regarding shikimate pathway flux, which means it was impossible to properly validate the entirety of the simulation ability of our model, particularly the shikimate pathway and the AAA production pathways.

### 2.3. Simulating AAA overexpressing mutant strains

Since there is a lack of quantitative experimental data on the shikimate and AAA pathways, we performed simulations with overexpressing strains found in the literature. In total, 41 different strains were selected, with 14 being specific to L-Phe production, 12 to L-Tyr production and 15 to L-Trp production. These strains were simulated using the dynamic model, using conditions similar to Chassagnole et al. However, not all characteristics of these strains genotype were able to be simulated, due to the model’s limitations. eing entirely based on kinetic mechanisms, characteristics such as gene repression or activation weren’t simulated. ikewise, due to the model’s size and structure, some of the reactions/genes found in the literature weren’t present in the model and thus, were impossible to be implemented into the simulation. Taking this into account all strains were successfully simulated using the dynamic model, with a single exception. This was most likely due to the number of KOs existing in that particular strain, which combined with the relatively small size of the dynamic model (lack of reactions and pathways to circumvent imposed conditions) led to a simulation failure. The relevant genotype of these strains can be seen in Table 2. One must note that, similar to the dynamic model, the stoichiometric model also couldn’t implement all of the strains genotype, most notably gene repression and activation. The stoichiometric model seems to have difficulty in simulating strains for L-Trp overexpression (OE), achieving no L-Trp production on any strain. Three of the L-Phe strains also show no OE in relation to WT simulation, while all L-Tyr overexpressing strains produce the expected phenotype. On the contrary, almost all strains successfully simulated with the dynamic model, with a single exception, present an OE of AAA when compared to its WT simulations, even L-Trp overexpressing strains. These results show a tendency for better prediction of AAA overexpression when using the dynamic model.

**Table 2.**
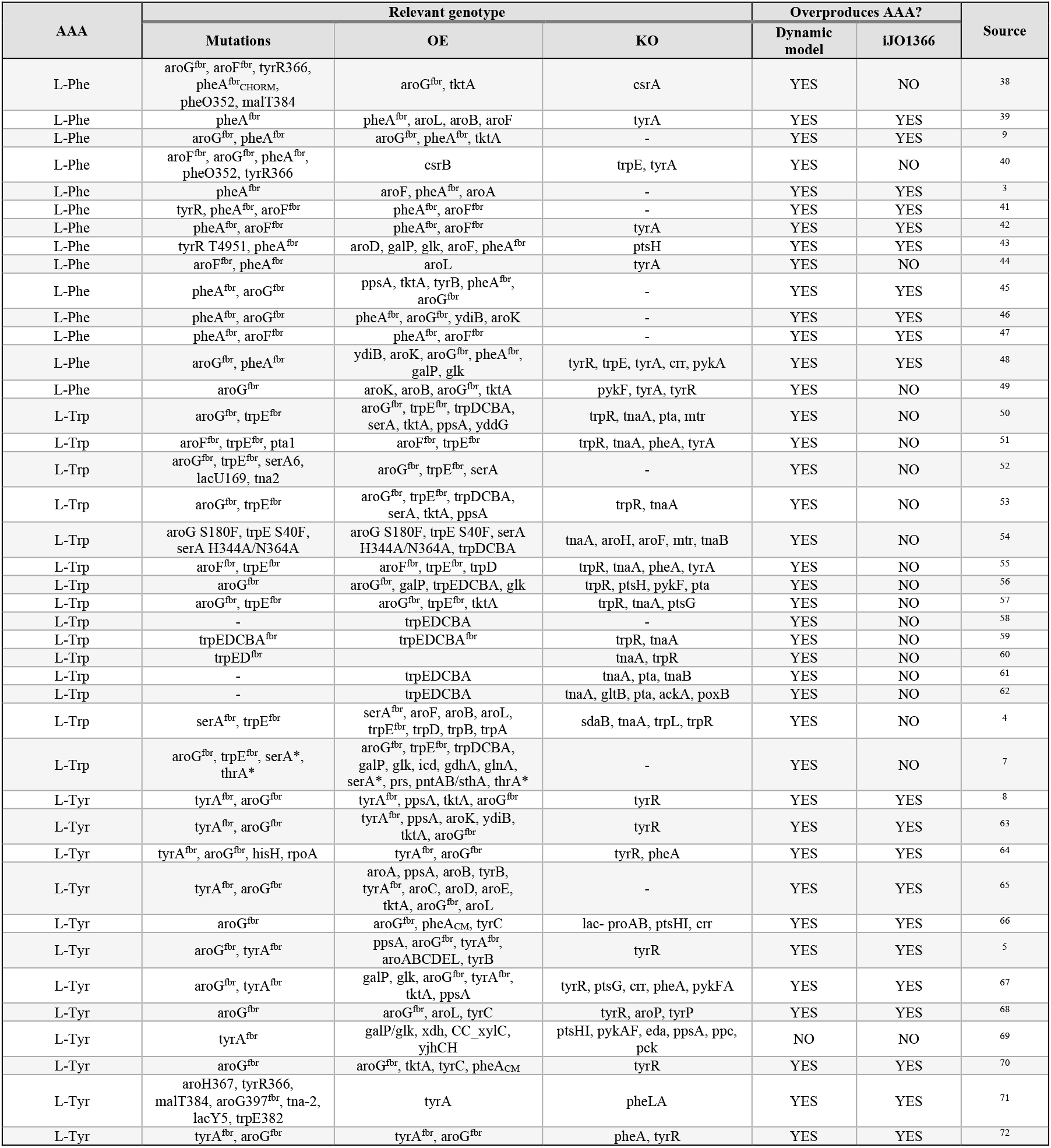
Strains for overexpression of AAA found in literature, with relevant genotype from each strain. Production of AAA in simulated strains is shown relative to WT, using dynamic and stoichiometric model (iJO1366). AAA – aromatic amino acids, OE – overexpression, KO – knockout, fbr – feedback inhibition resistance.

The literature strains presented in Table 2 show a high variability in targets for strain design, but there are common key targets found in several of the literature strains, for each AAA. Strains for L-Phe production typically possess feedback resistance mutations in genes that encode reactions sensitive to end-product inhibition, such as aroF/aroG or pheA. Besides these mutations, these genes are usually overexpressed. Then, to prevent carbon flow from diverting into the other AAA pathways, tyrA and trpE can be targets for KO.

Strains for L-Tyr production are quite similar to L-Phe strains, as they only differ in the two final steps of their production. The similarities include feedback resistance mutations in genes that encode reactions sensitive to end-product inhibition, such as aroG and tyrA, while also overexpressing these genes. To prevent carbon flow to divert into other pathways, pheA can be a target for KO. For both L-Phe and L-Tyr overexpressing strains, the removal of tyrR, a repressor, via mutation or KO is also a common strategy.

Strains for L-Trp OE are quite different from the other two as this amino acid is more tightly regulated, due to its longer pathway and requirement of additional compounds for its biosynthesis. These strains typically possess feedback resistance mutations in genes that encode reactions sensitive to end-product inhibition such as aroG/aroF or trpE. These genes are typically overexpressed. The entire operon for L-Trp biosynthesis is also typically overexpressed. Another two common targets are the KO of tnaA and trpR, which are responsible for tryptophan consumption and gene repression, respectively.

The dynamic model has been successful in simulating both gene KO and AAA overexpressing strains. However, it remains to be seen if it can be used as a target predictor for strain optimization studies. To test this, three rounds of optimization were performed with the dynamic model for each objective function, which was either L-Phe, L-Trp or L-Tyr intracellular concentration after 72000 s of simulation with the settings mentioned in Methods. The optimization solutions are analyzed in the following subsections.

### 2.4. Optimization results for Phenylalanine

In the *in silico* L-Phe optimization solutions, OE of tyrB and aroB are found in 100% of possible strains, despite having a much smaller presence in strains from literature (approximately 10%). This contrast could imply an overestimation of these genes’ importance by the model, but it could also mean that most reported strains have overlooked important targets. The gene pheA is another important target found in the *in silico* solutions, with over 90% of possible strains having OE and/or feedback resistance mutations of these gene. This value resembles the one from literature strains, over 90% of which also possess pheA^fbr^. However, pheA OE is only present in approximately 70% of literature strains. In the *in silico* solutions, other common targets include increasing inhibition sensitivity of trpE, OE of aroC and tyrA (chorismate mutase only), KO of glnA, and underexpression (UE) of glnA, pykAF, which are found in 35% to 55% of the possible strains. Increasing inhibition sensitivity of trpE can be substituted with trpE KO or UE, for equivalent results *in silico*. In fact, a small amount of literature strains have trpE KO as target. Surprisingly, OE of aroC and tyrA (chorismate mutase only) are not found in the literature strains, same as with KO and UE of glnA, making them new possible targets to create new strains of L-Phe overproduction. The *in silico* solutions also showed as target the KO or UE of one hypothetical gene, named here as geneA, which encodes a glutamate drain reaction in the model. Since the drain reaction is meant to represent a large number of metabolic fluxes, there is no clear answer to what this target is. We can only presume that removing reactions that degrade glutamate would be beneficial to increasing L-Phe production, assuming that removing one or several of those reactions is biologically viable.

The best *in silico* optimization solution has a higher L-Phe concentration than the simulated WT by a ratio of almost 5 times. This new possible strain’s genotype has targets already reported in literature and new possible targets. The latter include KO of genes glnA, gpsA and the hypothetical geneA, OE of genes aroC and gdhA, and UE of genes pykAF (which have been reported as KO targets, but never as UE; could be an alternative due to the importance of this gene). This solution can be found in Table 3. Other possible new strains and targets for L-Phe overproduction can be found annexed to this paper.

**Table 3.** Best strain optimization results for each of the three AAA. AA – amino acid, KO – knockout; OE – overexpression; UE – underexpression; fbr – feedback resistance; +inhib – increase enzyme sensitivity to inhibition by end product; gene targets in *italic* represent hypothetical genes; gene targets in **bold** represent targets not reported in the literature; other gene targets are reported in literature. Hypothetical genes are explained in text.

| AA | MUTATIONS | KO | OE | UE | x HIGHER THAN WT |
| --- | --- | --- | --- | --- | --- |
| L-PHE | *pheA<sup>fbr</sup>* | **glnA**, **gpsA**, *geneA* | *tyrB*, *aroB*, **aroC**, *pheA*, **gdhA** | **pfkAF** | 4.92 |
| L-TRP | *trpD<sup>fbr</sup>* | *tnaA*, *pheA*, **pck** | *trpED*, *prs*, *geneB* | **tyrA**, *geneA* | 1364 |
| L-TYR | **pheA<sup>+inhib</sup>**, *tyrA<sup>fbr</sup>* | **glnA**, *geneA* | *aroB*, *aroC*, *tyrA*, *tyrB*, *geneC* | **pheA** | 16.52 |

### 2.5. Optimization results for Tryptophan

In the *in silico* L-Trp optimization solutions there are four targets which are found in all of the best possible strains: KO of tnaA gene, OE of trpD and trpE and mutation of trpD to give it feedback resistance. These solutions are supported by literature, with these targets being present in over 50% of reported L-Trp overproducing strains, implying good prediction accuracy by the dynamic model. Other common targets also previously reported in literature strains include OE of prs and KO of pheA. In these *in silico* solutions there are also several new possible targets. One of them is UE of the gene tyrA, which has been reported in literature as KO. More data would be required to ascertain the differences between the two methods in an L-Trp overproducing strain. Other new possible targets include KO of tktB and pck and increased inhibition sensitivity of tyrA. KO of pck has been reported before, but L-Tyr overproducing strains. One can assume that due to the shikimate pathway being a common pathway to all AAA, KO of this gene could be a valid strategy for L-Trp strains. The *in silico* solutions also included increasing inhibition sensitivity of aroF, as well as UE of aroG and aroF. These seem counter intuitive as these genes encode the first reaction of the shikimate pathway and doing so should decrease carbon flow into the shikimate pathway. Experimental data would be required to evaluate these targets. The L-Trp optimization solutions also present hypothetical genes as targets, such as geneA (mentioned above), geneB and geneD. GeneB encodes a serine drain reaction, while geneD encodes a pyruvate drain reaction that represents synthesis of isoleucine, alanine, α-ketoisovalerate and diaminopimelate. Same as with geneA, we can presume that removing reactions that degrade serine and pyruvate (particularly for the synthesis of the mentioned compounds), could increase L-Trp production, assuming that removing those reactions is biologically viable.

The best *in silico* optimization solution has a higher L-Trp concentration than the simulated WT by a ratio of over 1000 times. This value is several times higher than for the other AAA, which could be due to the more constrained biosynthesis process of L-rp. his new possible strain’s genotype has targets already reported in literature, such as KO of tnaA and pheA, OE of trpE, trpD and prs, and feedback resistance mutations for gene trpD; as well as targets not yet reported, such as KO of pck, OE of geneB and UE of tyrA and geneA. This solution can be found in Table 3. Other possible new strains and targets for L-Trp overproduction can be found annexed to this paper.

### 2.6. Optimization results for Tyrosine

In the *in silico* L-Tyr optimization solutions, OE of genes aroB and tyrB are present in 100% of possible strains despite having much smaller presence in reported strains (around 10%). The same was noticed in L-Phe overproducing strains, both *in silico* and reported, which is expected due to the similarity between these two AAA biosynthesis. The gene pheA is another important target obtained by the *in silico* solutions, with over 80% of possible strains having UE and/or increased inhibition sensitivity mutations. Although this gene has been reported in literature strains before, it was as target for KO, thus these two could be new alternatives. More data would be required to evaluate the feasibility of these changes in a biological setting. In the *in silico* solutions, other common targets include OE of tyrA and aroC, and feedback resistance mutations of gene tyrA, all of which have been reported previously in literature strains. There are also new possible targets found, which include OE of gltB, KO or UE of glnA, and UE of pykAF. The KO of gene pykAF has been reported in literature to have a positive effect on L-Tyr production, but there is no data on UE of the gene. The *in silico* solutions also showed as target the KO or UE of hypothetical geneA, similarly to potential strains for L-Phe overproduction (the gene is mentioned above).

The best *in silico* optimization solution has a higher L-Tyr concentration than the simulated WT by a ratio of over times. his new possible strain’s genotype includes targets already reported in literature such as the OE of aroB, aroC, tyrA and tyrB, and the feedback resistance mutation of gene tyrA, as well as targets not yet reported such as KO of geneA, OE of geneC, UE of pheA and increased inhibition sensitivity of gene pheA. The hypothetical gene C encodes a drain reaction for phosphoenolpyruvate that represents mureine synthesis. Similar to the other hypothetical genes we can presume that removing reactions that degrade phosphoenolpyruvate, such as the synthesis of mureine, could increase L-Tyr production, assuming that removing those reactions is biologically viable. This solution can be found in Table 3. Other possible new strains and targets for L-Tyr overproduction can be found annexed to this paper.

### 2.7. Optimization results using iJO1366

Strain target optimization was also performed using the stoichiometric model iJO1366, to compare the differences in obtained strains and targets via this method. Despite the difference in size between the two models, optimization with iJO1366 gives solutions with fewer number of targets. In fact, the solutions obtained present only two targets, one of which was common for all solutions of the three AAA. This common target is non-viable to implement *in vivo* as it represents energetic balance in the form of ATP maintenance. The remaining targets all differ for each AAA. In L-he’s case, the OE of pheA or of tyrB are a possibility, both of which are targets already described in the literature. In L-rp’s case, of tna is the only other target found, apart from the common one. his wasn’t expected since KO of tnaA has already been described in literature in several strains and seems counter intuitive to L-Trp production since the reaction associated with tnaA uses tryptophan as substrate. In L-yr’s case, the OE of aroD or of aroL are a possibility, both of which have been described in literature. From these results, it seems that strain target optimization for the three AAA is better when using this dynamic model, as its solutions provide more possible targets, which are also consistent with *in vivo* findings from literature.

## 3. Methods

### 3.1. Modeling

There are several validated dynamic models in literature that represent *E. coli’s* CCM and other important pathways. These were used as basis for the creation of our own model. The sugar transport system, glycolysis, the pentose phosphate pathway, and extracellular kinetics were derived from Chassagnole et al.^12^; the tricarboxylic acid cycle, glyoxylate shunt, and co-metabolites maintenance were derived from Peskov et al.^23^; the acetate cycle was derived from Kadir et al.^22^; the intake of ammonia into the cell, glutamine and glutamate cycle was derived from Bruggeman et al.^73^; the electron transport chain and synthesis of ATP were derived from Peercy et al.^74^, Korla et al.^75^, and Ederer et al.^76^. The kinetics of some reactions of these models were adapted according to Lima et al.^15^, in order to fix some discrepancies in the units.

The shikimate and the AAA production pathways kinetics were obtained from *in vitro* results found in BRENDA^77^, or from the literature if not found in the BRENDA database. Kinetic parameters were chosen based on data quantity (papers with whole enzyme kinetic data were prioritized to papers with single parameter studies), existence of kinetic mechanism and paper publication date. In rare cases, when no kinetic information was found, complex mathematical expressions were reduced to empirical equations. The model consists of a system of ODEs that take different forms depending on their location regarding the cell. Intracellular metabolites take the form of equation 1:

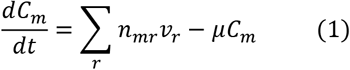

where *C* is the concentration of metabolite *m, t* is time, *µ* is the specific growth rate, *n*_*mr*_ is the stoichiometric coefficient for metabolite *m* in reaction *r*, the rate of which is *v*_*r*_. Extracellular substrates (glucose, acetate, and ammonia), take the form of equation 2:

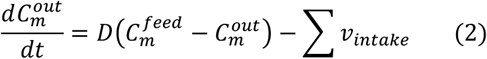

where C^out^ is the extracellular concentration, C^feed^ is the metabolite’s concentration in the feed, is dilution rate and v_intake_ is the intake reaction(s) for the extracellular metabolites.

The kinetics of all reactions, with mathematical equations and respective parameters, present in this model can be found annexed to this paper.

### 3.2. Simulation

All simulations and optimizations of the dynamic model were performed using optimModels (https://github.com/saragcorreia/optimModels), a Python package that allows simulation and strain optimization for dynamic models. OptimModels allows the solving of the system of ordinary differential equations that characterizes kinetic models. With this software, it is possible to simulate different phenotypes, change initial cellular conditions or change growth media. For simulation and optimization of the stoichiometric model, iJO1366, an open source software for simulation and metabolic engineering of stoichiometric models called Optflux^78^ was used. Model iJO1366 was simulated using flux balance analysis (FBA), with growth rate being set to experimental dilution rate values since, in continuous culture, at steady state, growth rate is equal to dilution rate.

With the dynamic model being based on enzyme kinetic information it doesn’t have data about gene-enzyme relationships. Therefore, simulation of strains with altered genome, such as single gene KO, was performed by studying the relationship between genes-enzymes in the literature and translating it into the model’s parameters. or instance, if a reaction follows the one gene one enzyme hypothesis, KO of a gene is equal to knocking out a reaction. However, if there is redundancy where a reaction is performed by two or more enzymes, which are encoded by two or more genes, one needs to consider the activity of each isoenzyme to know the direct effect that a particular gene KO would have on the reaction. With this in mind, to simulate different phenotypes/strains using the dynamic model kinetic parameters are altered, with the resulting phenotype being determined by the set of chosen parameters and alterations made. Changing the maximum reaction rates (Vmax) can mimic reaction KO, OE, or UE. Likewise, inhibition parameters (Ki) can also be altered, representing enzyme feedback resistance or even an increase in inhibition sensitivity by the enzyme. Michaelis-Menten constants (Km) can also be changed, implying a change in affinity for either substrates or products. These changes are done pre-simulation by increasing, decreasing, or nullifying these parameters by a factor of x (with x being a real number). To simplify explanation, number x will be referred to as Factor. A visual representation of this method is shown in Figure 2.

The model was prepared for simulation of a glucose pulse, similar to Chassagnole et al^12^. The simulations had a total time of 72000 s and a glucose initial concentration of 2 mM. The sole carbon source for this model is glucose. Acetate concentration in the feed and initial concentration were both 0 mM. The initial concentration of ammonia was 0.1 mM and its concentration in the feed was 14 mM. No other external metabolites are present in the model. Dilution rate and glucose concentration in the feed varied, according to the experimental conditions that were being simulated. While validating the model with experimental data by Ishii et al., the glucose concentration was set to 4 g/L and the dilution rate varied between 0.1 h^-1^ to 0.7 h^-1^. However, during the optimization tasks, these two parameters were set to 20 g/L and 0.1 h^-1^, respectively, similarly to Chassagnole et al.

The flux steady state values of both dynamic and stoichiometric models were normalized to the PTS (phosphotransferase) reaction. They were them compared to the experimental data by linear regression. The squared regression coefficient was calculated for each dilution rate and gene KO strain.

### 3.3. Optimization

All optimization tasks were performed with evolutionary algorithms (EA), which have been shown to be possible tools for strain design in metabolic engineering^19^.

EA are meta-heuristic approaches that simulate biological evolution processes, such as reproduction and mutation to obtain candidate solutions with better fitness values in regard to an objective function. The algorithm works by creating an initial population of different individuals, which are possible solutions with different characteristics. These individuals will be evaluated through an objective function and ranked according to their fitness, with the best ones being prioritized to pass on their characteristics. Different methods are applied to each individual (crossover or mutation), creating new individual candidates, which will create the next generation. Elitism is used to keep the best candidate from the previous generation in the new one. At the same time, newly created candidates will be added to the new generation to provide new characteristics, keeping the “gene pool” fresh. These steps are repeated until reaching a previously set final generation, with the goal of acquiring a better fitness value in each one, as represented in Figure 2. The characteristics for each individual were obtained by changing parameters, as mentioned above.

The Vmax and Ki parameters for all reactions were chosen as possible targets, in the exception of reactions related to co-metabolites, energy synthesis, feed intake, and drains that aren’t biologically viable (co-metabolite pathways, ETC and ATP synthesis). This was done to obtain solutions with variable characteristics (OE, UE, feedback resistance, KO, or inhibition sensitivity increase). The objective function used to evaluate each possible solution was the intracellular concentration of Phenylalanine, Tryptophan or Tyrosine. This function was chosen to create strains that overexpressed only one of the AAA, instead of all three. Each optimization task had two hundred generations, with a population of one hundred individuals per generation. The solutions had a possible size of up to ten targets (mutations, KOs, etc.). The solutions were filtered by fitness value, with only the solutions having over 90% of the highest fitness value found (in each optimization) being considered. The targets found in these solutions were then counted to find the most common ones, which are deemed more important by the optimization algorithm for AAA overexpression. The targets with over 10% participation in the best solutions were chosen for comparison with literature findings and stoichiometric model optimization results.

Strain target optimization using the stoichiometric model was performed with Optflux, using strength pareto evolutionary algorithms 2 (SPEA2). The objective function used was biomass-product coupled yield (BPCY), which takes the form of equation 3:

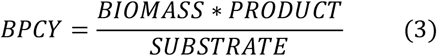

where BIOMASS is the simulated growth rate, PRODUCT is the exit reaction for the overproduced AAA, and SUBSTRATE is glucose intake. The simulation method chosen was FBA. A total of three optimization processes were run for each AAA, with 30000 solution evaluations (generations) per run. A maximum of 10 targets were allowed per solution, same as for the dynamic model.

## 4. Conclusion

The characteristics of the dynamic model presented in this paper allow for an enzyme kinetics point of view of *E. coli* metabolism. This in turn enables the use of details, which are usually discarded in other types of models, such as feedback inhibition. In fact, when using strain target optimization with this dynamic model, one can notice that feedback inhibition mutations are essential for an overproduction of the AAA. Several new possible targets and strains have been identified by strain target optimization with evolutionary algorithms, which allow for a non-biased approach to this problem. However, *in silico* simulations can only go so far and these new possible solutions require experimental validation. Overall, while this model still requires some quantitative validation, it has shown its great potential and the importance of this kind of models for strain target discovery, while at the same time providing new possible targets and strains for AAA overproduction.

